# Spontaneous emergence of azithromycin resistance in independent lineages of *Salmonella* Typhi in Northern India

**DOI:** 10.1101/2020.10.23.351957

**Authors:** Megan E. Carey, Ruby Jain, Mohammad Yousuf, Mailis Maes, Zoe A. Dyson, Trang Nguyen Hoang Thu, To Nguyen Thi Nguyen, Thanh Ho Ngoc Dan, Quynh Nhu Pham Nguyen, Jaspreet Mahindroo, Duy Thanh Pham, Kawaljeet Singh Sandha, Stephen Baker, Neelam Taneja

## Abstract

**Background:** The emergence and spread of antimicrobial resistance (AMR) pose a major threat to the effective treatment and control of typhoid fever. The ongoing outbreak of extensively drug resistant (XDR) *Salmonella* Typhi (*S*. Typhi) in Pakistan has left azithromycin as the only remaining broadly efficacious oral antimicrobial for typhoid in South Asia. Ominously, azithromycin resistant *S*. Typhi organisms have been subsequently reported in Bangladesh, Pakistan, and Nepal.

**Methods:** Here, we aimed to understand the molecular basis of AMR in 66 *S*. Typhi isolated in a cross-sectional study performed in a suburb of Chandigarh in Northern India using whole genome sequencing (WGS) and phylogenetic analysis.

**Results:** We identified seven *S*. Typhi organisms with the R717Q mutation in the *acrB* gene that was recently found to confer resistance to azithromycin in Bangladesh. Six out of the azithromycin-resistant *S*. Typhi isolates also exhibited triple mutations in *gyrA* (S83F and D87N) and *parC* (S80I) genes and were resistant to ciprofloxacin. These contemporary ciprofloxacin/azithromycin-resistant isolates were phylogenetically distinct from each other and from those reported from Bangladesh, Pakistan, and Nepal.

**Conclusions:** The independent emergence of azithromycin resistant typhoid in Northern India reflects an emerging broader problem across South Asia and illustrates the urgent need for the introduction of typhoid conjugate vaccines (TCVs) in the region.

**Key points:** We identified ciprofloxacin/azithromycin-resistant *Salmonella* Typhi (*S*. Typhi) in Chandigarh in Northern India. The independent emergence of ciprofloxacin/azithromycin-resistant typhoid in Bangladesh, Pakistan, Nepal, and India and the continued spread of extensively-drug resistant (XDR) typhoid in Pakistan highlight the limitations of licensed oral treatments for typhoid fever in South Asia.

## Background

*Salmonella enterica* serovar Typhi (*S*. Typhi), the etiologic agent of typhoid fever, is associated with an estimated 10.9 million infections and 116,800 deaths globally^1^. The majority of this disease burden is concentrated in South Asia, which has a modelled incidence rate of 592 cases per 100,000 person-years^1^. A pooled estimate of typhoid fever incidence of 377 cases per 100,000 in India has also been calculated using limited population-based data; significant geographical heterogeneity was observed^2^. The ongoing Surveillance for Enteric Fever in India (SEFI) study is generating geographically representative, age-specific incidence data, as well as additional information regarding cost of illness, range of clinical severity, and antimicrobial resistance (AMR) patterns associated with *S*. Typhi in India^3^. These data will undoubtedly provide a more comprehensive understanding of typhoid fever incidence rates across the Indian sub-continent, and ultimately, in supporting decision-making concerning typhoid conjugate vaccine (TCV) introduction in India ^4^.

Growing rates of AMR have made typhoid control increasingly challenging, beginning with the rise of multi-drug resistance (MDR; resistant to chloramphenicol, trimethoprim-sulfamethoxazole, ampicillin) in the 1990s,^5^ and the subsequent increase in fluoroquinolone resistance in the early 2000s, which was predominantly focused in South and Southeast Asia^6,7^. Ultimately, these phenotypic changes led to the common use of third generation cephalosporins for treatment of typhoid fever. The emergence and spread of extensively-drug resistant (XDR; resistant to chloramphenicol, ampicillin, cotrimoxazole, streptomycin, fluoroquinolones, and third-generation cephalosporins) typhoid in Pakistan has left azithromycin as the only available oral antimicrobial for effective treatment of typhoid fever across South Asia^8^. Concerningly, azithromycin-resistant *S*. Typhi have subsequently been reported in Bangladesh, Pakistan, and Nepal, although this phenotype has not, as yet, arisen in XDR organisms^9–11^.

It is apparent that we have an escalating problem with drug resistant *S*. Typhi in South Asia, due in part to empirical treatment of febrile patients and widespread community availability of antimicrobials. We are currently unsure of the regional distribution of azithromycin resistance or the potential for the emergence of a specific sub-lineage with this phenotype; as has been observed for MDR, XDR, and fluoroquinolone resistance. Here, we aimed to characterize the molecular basis of AMR in *S*. Typhi in a cross-sectional study performed in a suburb of Chandigarh in Northern India. Through whole genome sequencing (WGS), we describe the distribution of a collection of azithromycin resistant *S*. Typhi and show that these organisms have arisen independently of those in Pakistan and Bangladesh through the acquisition of an identical mutation in *arcB*. Our data support the prioritization of TCV introduction to prevent the continued emergence and spread of drug-resistant *S*. Typhi infections in South Asia.

## Methods

### Ethics

Ethical clearance was granted by the Institutional Ethics Committee of the Government Multi-specialty Hospital, Sector 16, Chandigarh (letter no GMSH/2018/8763 dated 26.7.2018) and the Postgraduate Institute of Medical Education & Research (PGIMER) Institutional Ethics Committee (IEC-08/2018-285 dated 24-9-2018). Administrative approval to carry out this collaborative study on enteric fever was granted by the Chandigarh Health Department (CHMM-2017/2991 dated 28.8.2017). Approval was also granted by the Collaborative Research Committee of PGIMER (no 79/227-Edu-18/4997 dated 12/12/2018). Informed consent was a prerequisite for inclusion in the study.

### Study design

The *S*. Typhi isolates were obtained from blood cultures taken from febrile patients presenting to Civil Hospital Manimajra in Chandigarh (CHMM) between September 2016 and December 2017, where passive blood culture surveillance has been conducted since November 2013. CHMM is a 100-bed secondary health care facility located on the outskirts of Chandigarh, and serves a catchment area of approximately 200,000 people, including referrals from four primary care centers. Data from patients with blood culture confirmed invasive *Salmonella* infection, which includes *Salmonella* serovars Typhi and Paratyphi A, B and C, from both inpatient and outpatient wards were used in the study. Clinical history, laboratory test results, and risk factor data were recorded for each confirmed case of enteric fever for patients residing in Manimajra. This analysis focuses solely on confirmed cases of *S*. Typhi.

### Identification and Antimicrobial Susceptibility Testing

Bacterial isolates were identified as *S*. Typhi using conventional biochemical tests; Motility agar, Hugh–Leifson Oxidative-Fermentation (OF) test, the Triple Sugar Iron (TSI) test, citrate test, urease test, phenyl pyruvic acid (PPA) test, and indole test. All isolates were eventually confirmed using antisera from Central Research Institute (CRI), Kasauli. Antimicrobial susceptibility was determined for the following antimicrobials by disc diffusion: ampicillin (10μg), chloramphenicol (30μg), trimethoprim/sulfamethoxazole (1.25/23.75μg), ceftriaxone (30μg), azithromycin (15μg), ciprofloxacin (5μg), and pefloxacin (5 μg). Zone diameters were measured and interpreted as per Clinical Laboratory Standards Institute (CLSI) guidelines^12^. Minimum inhibitory concentration testing was also conducted on all organisms showing resistance to any of the above antimicrobials by disc diffusion using E-tests (bioMerieux, France).

### Whole-Genome Sequencing and Phylogenetic Analysis

All *S*. Typhi were stored and shipped to PGIMER Chandigarh. Isolates from patients residing outside of Manimajra were excluded, as were those for which there was inadequate clinical metadata, or the DNA yield was below the amount required for WGS. Total genomic DNA was extracted from the *S*. Typhi using the Wizard genomic DNA extraction Kit (Promega, Wisconsin, USA) and subjected to WGS using the Illumina MiSeq platform (Illumina, San Diego, CA, USA) to generate 250 bp paired end reads. We then aimed to put these sequences into global and regional context. These reads were mapped against the CT18 reference sequence (accession no. AL513382) using the RedDog mapping pipeline (available at: https://github.com/katholt/RedDog) to identify single nucleotide variants (SNVs) ^13–15^. After removing prophages and recombinant^16^ and repetitive sequences, we generated a final alignment of 25,832 chromosomal SNVs for 3472 isolates. SRST2^17^ was used with ARGannot^18^ and PlasmidFinder^19^ to identify AMR genes and plasmid replicons, respectively. Mutations in *gyrA*, and *parC*, as well as the R717Q mutation in *acrB* were detected using GenoTyphi (https://github.com/katholt/genotyphi). Maximum likelihood phylogenetic trees were inferred from the chromosomal SNV alignments using RAxML (v8.2.9)^20^, and then visualized in Microreact^21^ (https://microreact.org/project/nniNzBL2uq3XZXYDKgG374) and the Interactive Tree of Life (ITOL)^22^. Raw read data were deposited in the European Nucleotide under accession number ERP124488.

## Results

### Epidemiological observations

Typhoid fever is a major public health concern among children and young adults in this region of Northern India. It is thought that many cases in this area, a hub for the states of Punjab, Haryana, and Himachal Pradesh, are associated with the mixing of large populations of seasonal workers from the adjoining states. These workers generally live in informal dwellings with poor sanitation and limited access to safe water. Approximately 1,500 suspected enteric fever patients present to the CHMM facility annually and receive blood cultures, of which ~10% are positive for *S*. Typhi. There is a seasonal peak of typhoid fever in this facility during the monsoon months from May to September (Figure 1).

**Figure 1.**
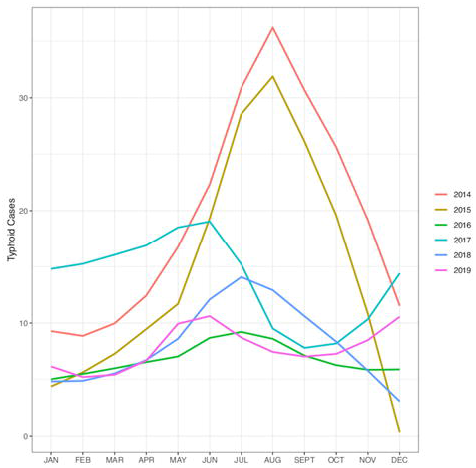
The annual seasonality of typhoid fever in Manimajra, Chandigarh. Plots showing the number of typhoid cases recorded the civil hospital in Manimajra from 2014 to 2019. The observed annual peak in typhoid cases corresponds with the monsoon season in Northern India (May to September).

The *S*. Typhi organisms interrogated here by WGS were isolated between September 2016 and December 2017 and all originated from blood cultures taken from febrile patients attending CHMM. All patients with a positive blood culture for *S*. Typhi resided within 12.9 km of the healthcare facility and were located in an area of ~28 km^2^ (Figure 2). *S*. Typhi was isolated throughout the specified months, again with higher number of cases observed between May and September. The median age of typhoid patients included in this analysis was seven years. The standard of care antimicrobials at this facility for patients with suspected enteric fever in outpatient settings are cotrimoxazole, cefixime, and/or azithromycin, and ceftriaxone for inpatients.

**Figure 2.**
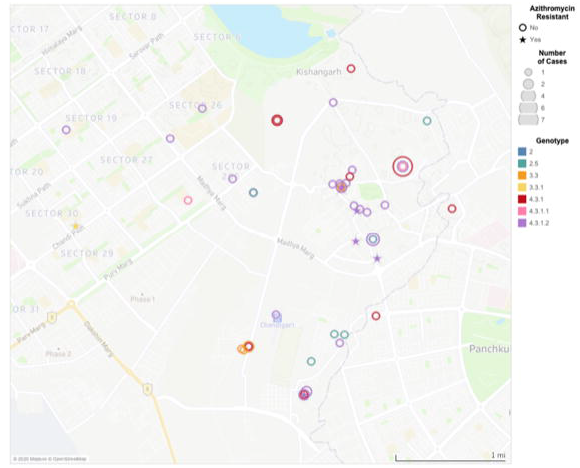
The spatial distribution of confirmed typhoid fever cases in Chandigarh. Map of Chandigarh (scale shown) of the residential locations of the *S*. Typhi cases recorded at the civil hospital in Manimajra between September 2016 and December 2017. *S*. Typhi genotype is indicated by color (see key), azithromycin-resistant isolates are indicated with stars, and the number of cases in each coordinate is represented by the size of the circles. All cases were located within a 28 km^2^ area. There is a cluster of cases of genotype 4.3.1.2 in a 0.25 km^2^ area of central Manimajra, which includes five of the six closely related azithromycin-resistant isolates.

### The local phylogenetic structure of S. Typhi

Ultimately, after data quality control, we generated and analyzed 66 *S*. Typhi genome sequences from Chandigarh. We observed that the population structure of *S*. Typhi around Chandigarh exhibited a high level of genetic diversity with eight co-circulating genotypes, indicative of population mixing and sustained introduction of organisms from a variety of locations across India (Figure 3). However, and in an analogous manner to other locations in Asia (and East Africa), most organisms (80%; 53/66) belonged to lineage 4.3.1 (H58), with the majority of those (66%; 35/53) belonging to 4.3.1.2. In total, 24% (16/66) of isolates were subclade 4.3.1, and 3% (2/66) were 4.3.1.1. Additional genotypes were comprised of subclade 3.3 (7.5%, 5/66), clade 2.5 (7.5%, 5/66), clade 3.3.1 (1.5%, 1/66), clade 4.1 (1.5%, 1/66) and major lineage 2 (genotype 2; 1.5%, 1/66).

**Figure 3.**
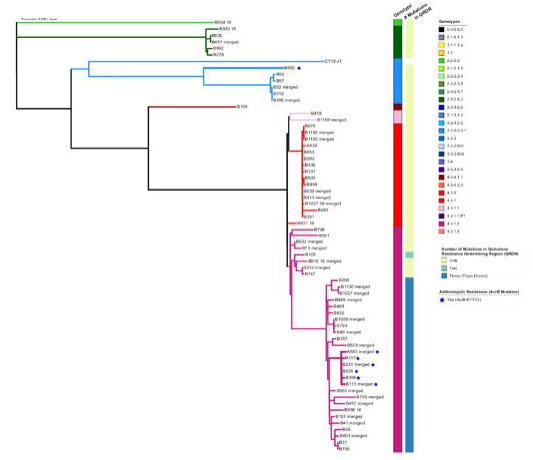
The phylogenetic distribution of *S*. Typhi isolated at the civil hospital in Manimajra, Chandigarh. Phylogenetic tree made in Ram of the 67 isolates genome sequenced. This collection shows considerable genetic diversity, with eight genotypes represented (as colour coded on branches and in the key). Mutations in the Quinolone Resistance Determining Region (QRDR), and presence of the acrB-R717Q mutation are shown for each organism. There are two distinct clusters of organisms with the *acrB* mutation that confers azithromycin resistance; each of these individual organisms indicated with a star.

### Fluoroquinolone resistance

All (66/66) *S*. Typhi genome sequences, regardless of the genotype, possessed mutations in *gyrA*, conferring reduced susceptibility to fluoroquinolones. Notably, given that these mutations were observed in a range of genotypes, these had occurred independently, likely as a result of sustained antimicrobial pressure from widespread fluoroquinolone use. We further observed multiple *gyrA* mutation profiles in 4.3.1 organisms conferring intermediate resistance against fluoroquinolones (0.12μg/ml < ciprofloxacin MIC < 1μg/ml). These mutations included S83Y (29.1%; 16/55), S83F (16.4%; 9/55), and D87N (1.8%, 1/55). Additionally, we identified a subclade of organisms that represented 49.1% (27/55) of the 4.3.1 isolates, all of which belonged to 4.3.1.2, that contained the classical triple mutations associated with fluoroquinolone resistance (S83F and D87N in *gyrA* and S80I in *parC*)^14^. These organisms exhibited high-level fluoroquinolone resistance (ciprofloxacin MIC >24μg/ml). Our observations with respect to ubiquitous fluoroquinolone resistance were concerning; however, none of the *S*. Typhi isolates were MDR, which may be associated with a reduced reliance on older classes of antimicrobials.

### Azithromycin resistance

We identified that 7/66 (10.6%) of the sequenced isolates contained a mutation in *acrB*, a gene encoding a component of the AcrAB efflux pump^23^. Mutations in *acrB* have been previously observed to be associated with resistance to azithromycin^9^. Here, the *acrB* mutation was non-synonymous (R717Q) and identified in six genotype 4.3.1.2 organisms and in one genotype 3.3.1 organism. These data are indicative of convergent mutation in different lineages, highlighting a potential increasing reliance on azithromycin; this selective pressure is further accentuated by the small clonal expansion in genotype 4.3.1.2 (Figure 3).

The R717Q mutation in *acrB* has been linked to high azithromycin MICs in genotype 4.3.1.1 *S*. Typhi isolates from Bangladesh^8^ and also in genotype 4.3.1.1 *S*. Typhi in Pakistan^9^. The contemporary Indian *S*. Typhi isolates identified here with the same R717Q mutation in *acrB* showed resistance to azithromycin with an MIC >16μg/ml (range: 16 ->256μg/ml). Our data suggested that these R7171Q mutations in the *acrB* gene have arisen spontaneously in India. To test this hypothesis, we constructed an expanded phylogenetic tree comprising a global *S*. Typhi collection, including organisms from across South Asia and the recently described azithromycin-resistant organisms from Bangladesh, Pakistan, and Nepal. We found that the azithromycin-resistant *S*. Typhi from India were phylogenetically distinct from those reported from Bangladesh, Pakistan, and Nepal (Figure 4). Additionally, we found that the azithromycin-resistant organisms associated with *acrB* mutations were dispersed around the tree and appear to have arisen on at least five different occasions, with a differing *acrB* mutation in organisms from Nepal (R717L)^11^.

**Figure 4.**
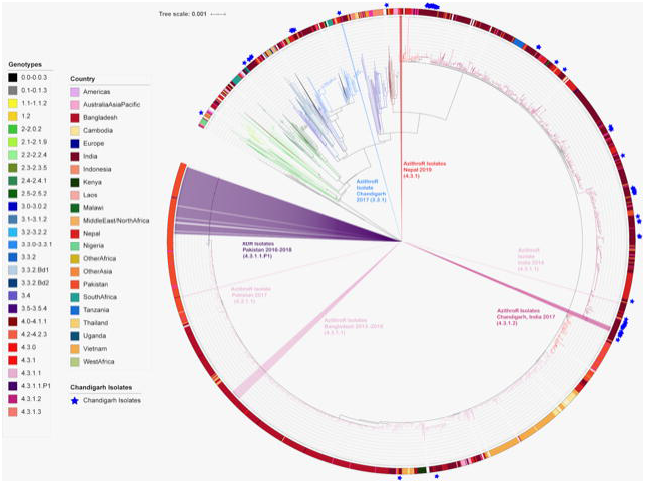
Azithromycin resistant *S*. Typhi in global context. Diagram depicts a maximum likelihood rooted phylogenetic tree with a final alignment of 25,832 chromosomal SNVs for 3,472 globally representative isolates, including all publicly available isolates from India. The colour of the internal branches represents the genotype, the colored ring around the tree indicates the country or region of origin for each isolate, and the blue stars indicate which isolates were originate from this study. Additionally, the tree contains each known *S*. Typhi isolate with an *acrB* mutation in public databases, these originate from India, Nepal, Bangladesh, and Pakistan. The location of the XDR isolates from Pakistan are added for context.

The azithromycin-resistant isolates from India described here were isolated in 2017, meaning that they are contemporaneous with those reported from Bangladesh and Pakistan, and arose independently in phylogenetically distinct lineages. Lastly, six of the seven Indian isolates with the R717Q mutation in *arcB* (all 4.3.1.2) were also within the group of organisms with the triple mutation associated with high level fluoroquinolone resistance, making these organisms highly resistant to these two key oral antimicrobials.

## Discussion

In this study, we aimed to describe the genomic aspects the *S*. Typhi causing disease in an endemic region in Northern India. We investigated antimicrobial susceptibility patterns using phenotypic testing and WGS data and then placed these data into a regional and global context using published genomic data. Notably, we identified seven azithromycin resistant *S*. Typhi isolates. These organisms belonged to two different lineages and were genetically distinct from azithromycin resistant isolates recently reported from Bangladesh, Pakistan, and Nepal. Our observations suggest azithromycin resistance mutations at codon 717 in *arcB* are occurring independently in locations where there is substantial selective pressure induced by azithromycin. An increased reliance on azithromycin for treatment of typhoid fever and other invasive bacterial infections in South Asia, along with ongoing clinical trials measuring the impact of prophylactic administration of azithromycin on growth and mortality of infants and young children in Pakistan, Bangladesh, and India,^24^ signal the inevitability of what Hooda and colleagues have termed pan-oral drug-resistant (PoDR) Typhi^24^. This scenario would necessitate inpatient intravenous drug administration for effective treatment of typhoid fever in the region at enormous cost to patients and to healthcare systems. Where intravenous drug administration is not an option, typhoid could once again become a disease with a high mortality rate, as was observed in the pre-antimicrobial era.

While the catchment area of this study is not representative of the entire Indian sub-continent, the phenomenon described herein is unlikely to be restricted to Chandigarh. India is currently the largest consumer of antimicrobials of all LMICs, with a reported 6.5 billion Defined Daily Doses (DDDs) in 2015, or 13.6 DDDs per 1,000 inhabitants per day^25,26^. With such widespread availability and use of antimicrobials nationally, selective pressure on circulating pathogens is likely immense. The SEFI study will soon yield additional concrete AMR data from multiple, geographically representative sites in India, which will further elucidate AMR patterns across the country. Ideally, these data will inform local antimicrobial stewardship practices and may be used as a basis for prioritization of future interventions.

How then to tackle this red queen dilemma? Drug development efforts cannot keep pace with bacterial evolution. Therefore, there is an urgent need for preventative interventions, namely water, sanitation and hygiene (WASH) interventions and TCV introduction, in India and across South Asia. There is also a need for enhanced typhoid surveillance, in South Asia and globally, specifically to monitor the emergence and spread of this and other resistance phenotypes. Historic genomic data show us that drug-resistant *S*. Typhi lineages emerge in South Asia and then spread to East Africa and even Latin America^27^. With the widespread prophylactic deployment of azithromycin through clinical studies and public health programs in West Africa and South Asia,^24,28^ it will be critical to monitor global AMR patterns to mitigate a public health catastrophe.

This emerging problem additionally represents an opportunity for use of genomics to inform policy. WGS data provides clear information regarding AMR in organisms where molecular mechanisms of resistance are understood. Genomic surveillance also enables the identification and characterization of new resistance phenotypes, as was the case for XDR typhoid^8^ and the new azithromycin resistant organisms identified in Bangladesh and described further here^9^. The outputs of antimicrobial susceptibility testing are not always straightforward, particularly in cases where susceptibility breakpoints have not been validated extensively using clinical data, as is the case for azithromycin^29^. Genomic AMR data can inform prioritization of TCV introduction, as well as implementation of WASH interventions. Genomic surveillance should also be an important component of long-term monitoring of the impact of widespread TCV deployment. Not only can genomic surveillance provide additional information on the impact of TCV on AMR, it will also illustrate the impact of vaccine on bacterial population structures and enable the identification of any vaccine escape mutants. Such information is vital to understanding the long-term impact of vaccine, and to facilitate any potential future efforts for global typhoid elimination.

## Acknowledgements

We would like to thank the patients in Manimajra who participated in the study and the health care workers who screened and enrolled patients.

